# Molecular diet analysis in zebra and quagga mussels (*Dreissena* spp.) and an assessment of the utility of aquatic filter feeders as biological eDNA filters

**DOI:** 10.1101/2021.03.01.432951

**Authors:** Sven Weber, Lukas Brink, Manuel Wörner, Sven Künzel, Michael Veith, Diana Teubner, Roland Klein, Martin Paulus, Henrik Krehenwinkel

## Abstract

Molecular gut content analysis is a popular tool to study food web interactions and was recently also suggested as an alternative source for DNA based biomonitoring. However, the overabundant consumer’s DNA often outcompetes that of its diet during PCR. Blocking approaches are an efficient means to reduce consumer amplification while retaining broad specificity for dietary taxa. We here designed an assay to monitor the eukaryotic diet of mussels and test their utility as biological eDNA filters to monitor planktonic communities. We designed several rDNA primer sets with a broad taxonomic suitability for eukaryotes, which suppress the amplification of mussels. The primers were tested using mussel DNA extracts and the results were compared to eDNA water samples collected next to the mussel colonies. Taxonomic recovery, as well as patterns of alpha and beta diversity, were compared between mussels and water samples. In addition, we analyzed time series samples of mussel samples from different German rivers. Our primer sets efficiently block the amplification of various mussel genera. The recovered DNA reflects a broad dietary preference across the eukaryotic tree of life and considerable taxonomic overlap with filtered water samples. We also recover various taxa of possible commensals and parasites, associated with the mussels. Our protocol will enable large scale dietary analysis in mussels, facilitate aquatic food web analysis, elucidate the ecological impact of invasive bivalves and the rapid survey of mussel aquacultures for pathogens. Moreover, we show that mussels could serve as an interesting complementary DNA source for biomonitoring.

## Introduction

The immense task of documenting human imprints on ecosystems has been greatly simplified by eDNA metabarcoding (Taberlet et al. 2012). The filtering of water and sequencing of the filtrate is a commonly used approach in aquatic eDNA analysis (Barnes et al. 2020), but various other substrates can serve as eDNA sources. Examples include sediments, marine cobbles or the gut content of detritus feeding animals (Koziol et al. 2019; Shum et al. 2019; Siegenthaler et al. 2019). Another promising source of aquatic eDNA are sponges (Mariani et al. 2019), which behave like a biological eDNA filter, filtering and retaining eDNA particles from their surrounding community.

Besides sponges, a particularly well-suited organism to target as a biological eDNA filter are mussels. Bivalves are ubiquitous inhabitants of aquatic environments, important components of most limnic and marine food webs (Vaughn et al. 2008; Newell 2004) and of great economic importance in aquaculture (Shumway et al. 2003). Considering their ecological and economic importance, the efficient characterization of the natural diet of mussels is also of critical importance for aquatic food web analysis, to understand their impact on ecosystems and to optimize and survey aquaculture.

Mussels filter the water column for plankton and detritus (Lavrentyev et al. 1995; MacIsaac et al. 1995; Wong & Levinton 2004). The filtering mechanism is highly efficient: invasive mussels, which are known to build up high densities within a short time, rapidly alter plankton composition, leading to state shifts in entire ecosystems (Maguire & Grey 2006; Miller & Watzin 2007). While mussels can show certain diet selectivity (Baker & Levinton 2003), their filtering mechanism opportunistically retains most particles of a certain size range (Sprung & Rose 1988). The size of particles retained by mussels is well within that of common eDNA water filters (0.2-10 µm) (Barnes et al. 2020; Wilcox et al. 2015).

So far, diet analysis in mussels is mostly based on laboratory feeding assays or chemical screens (Kreeger et al. 1993;Petterson et al. 2010; Fernández et al. 2015). Metabarcoding approaches now offer a powerful alternative. To characterize a mussel’s diet, consumed DNA could simply be amplified from DNA extracts of the mussel’s gill and intestinal tissue. Universal PCR primers could be used to enable the recovery of a broad range of dietary taxa. However, with such universal primers, the highly overabundant consumer DNA may outcompete dietary taxa during PCR (Krehenwinkel et al. 2017). A pragmatic solution to this problem is the use of very high sequencing coverage. Consumer sequences are removed from the data before analysis (Pinol et al. 2015). However, the DNA of dominant taxa is often so overabundant that nearly no desired sequences remain (Krehenwinkel et al. 2017). An alternative solution is the use of blocking approaches, which prevent the consumer’s DNA from being amplified (Vestheim & Jarman 2008). Diagnostic SNPs at the primer’s 3’-end are highly efficient at blocking amplification of target lineages (Stadhouders et al. 2010; Krehenwinkel et al. 2019). This allows the recovery of even minute amounts of dietary DNA. Suitable PCR primers can be designed in conserved sequences, assuring a broad taxonomic specificity.

Here, we aim to: 1) develop a metabarcoding assay that allows the reconstruction of mussel dietary composition across the whole eukaryotic tree of life, while at the same time excluding mussels from amplification by 3’-primer mismatches. And 2) determine whether this assay provides a suitable alternative or complement to aquatic eDNA analysis of eukaryotic communities from filtered water samples.

We particularly focus on two species of the genus *Dreissena*, the zebra mussel *D. polymorpha* and the quagga mussel *D. rostriformis*. Both species’ native range is in far Eastern Europe, but they are now widespread invasives in Western Europe and America (Kinzelbach 1992; Son 2007; Heiler et al. 2013; Nalepa & Schloesser 2013; Paulus et al. 2014), where they cause great ecological and economic damage (Maguire & Grey 2006; Connelly et al. 2007; Miller & Watzin 2007). Considering their great impact, a protocol for the detailed assessment of dietary preferences of *Dreissena* spp. is particularly relevant.

We designed several primer pairs targeting the nuclear 18S ribosomal DNA. These primers were located in highly conserved sequences across the eukaryote tree of life to guarantee a broad taxonomic coverage. At the same time, they encompassed highly variable loop sequences, which allowed to discriminate closely related taxa. One set of primers efficiently suppressed amplification of *Dreissena* and mussels of 34 other genera in 16 families, due to two lineage diagnostic SNPs. Another set was designed to reduce amplification of metazoans and recover non metazoan dietary taxa from any filter feeding metazoan. We tested our primers on different samples of zebra and quagga mussels. In parallel, we collected several eDNA water samples in close proximity to the sampled mussel colonies, which were directly compared to the mussel samples. To test the broad applicability of our assay, we also included samples of the marine blue mussel *Mytilus edulis*. To show a direct application of our protocol, we analyzed several time series samples of *D. polymorpha*, collected in three German rivers over the past 25 years by the German Environmental Specimen Bank (hereafter ESB). The ESB collects these mussels annually as a bioindicator for water pollution following the guideline of Teubner et al. (2018), which guarantees comparability of samples among sites and across years. Using our protocol with ESB samples, we show differences of prey communities between rivers and temporal changes of mussel associated eDNA.

## Methods

### Design of blocking primers to recover the diet of mussels

Mitochondrial COI is the marker of choice for metabarcoding in the majority of metazoan groups (Elbrecht & Leese 2017, Marquina et al. 2019, Thomsen & Sigsgaard 2019). Mussels’ diet, however, consists of taxonomically diverse eukaryotic plankton, including different plant and algal groups, various protozoans and metazoans. COI is not a well-established barcode marker for many of these groups. A more suitable universal target locus for these taxa is found in the nuclear ribosomal RNA genes (Cabra et al. 2016, Giebner et al. 2020, Seymour et al. 2020). Here we focused on the 18SrDNA as a potential marker to exclude *Dreissena* from amplification, while retaining a broad specificity for various planktonic taxa. We generated an alignment of 158 near complete sequences of the 18S gene for different eukaryotic taxa (Genbank, assessed September 2020, see supplementary material) and screened this region for possible priming sites that would suppress the amplification of *Dreissena* mussels, while at the same time retaining a broad specificity for the remaining eukaryotes. A particularly suitable region for our purpose was found in the variable V8 and V9 regions of the gene, which are already widely used to generate barcoding markers for eukaryotes (Machida & Knowlton 2012, Albaina et al 2016, Choi & Park 2020, Sidker et al. 2020). A highly conserved fragment at the 5’-end of the gene was used to design reverse primers. In comparison to other eukaryotes, *Dreissena* spp. show two diagnostic substitutions from AA to TC at this position. This results in an A-A and an A-G mismatch at the last two positions of the primer, leading to pronounced drop in amplification efficiency of *Dreissena* (Stadhouders et al. 2010).

To explore the generality of the observed patterns of mismatches, we downloaded the exact mismatch sequence of more than 10,000 genera across the eukaryotic tree of life (See supplementary material). Only one random member per genus was included in the analysis, to avoid biases by species rich genera, or groups where many sequences of single species were deposited. Out of all analyzed eukaryotic genera, only 47 (0.42 %) showed the *Dreissena* blocking sequence (3’-TC) (Figure 1 b). 34 of these genera were bivalves (in 16 families), 8 belonged to other metazoans (5 annelids, 1 arthropod and one gastrotrich), and the remaining 5 were fungi and one oomycete.

**Figure 1.**
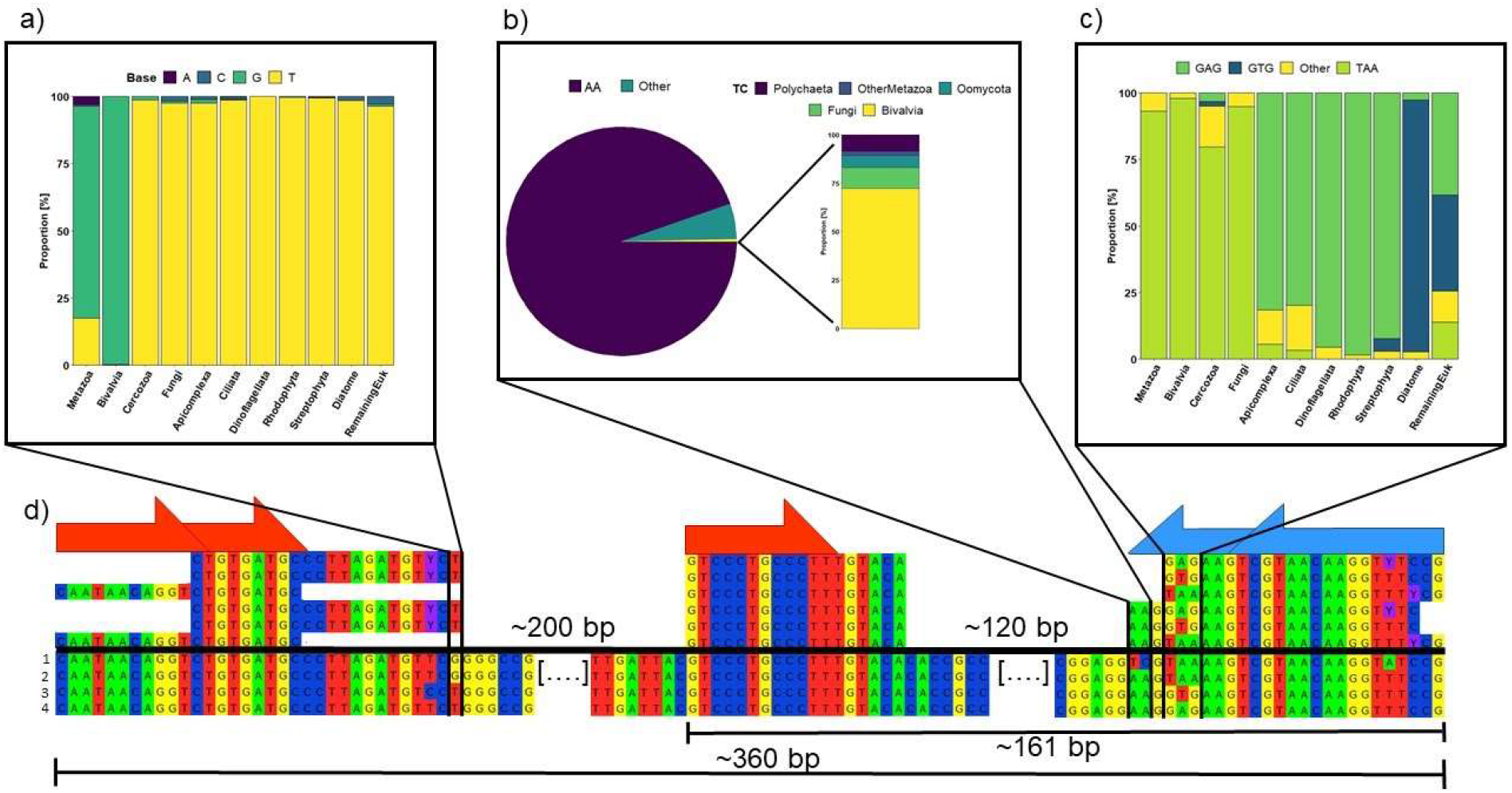
Overview of the designed primer combinations and their mismatches with different eukaryotic groups. The lower alignment shows subsections of the amplified fragment, highlighting the amplified fragment lengths and primer binding sites for a (1) Dreissena and (2) a Mytilus mussel, followed by (3) a brown algae and (4) a green algae. The upper six sequences show the different primer combinations we designed. The taxon specific mismatches are highlighted by black bars. The insets show the specificity of each mismatch across a total of ∼10,000 different genera from various eukaryotic lineages. shows the distribution of a diagnostic 3’-T or G across different eukaryotes. b) shows the distribution of 3’-AA vs. TC across eukaryotes. Only 0.42 % of all tested eukaryote genera show the Dreissena blocking sequence TC here as seen in the barplot magnified from the pie chart; and the majority of those genera are bivalves. c) shows the distribution of a diagnostic 3’-TAA, GAG or GTG across different eukaryotes.

In addition to blocking *Dreissena* from amplification, the fourth, fifth and sixth positions of the priming site contained substitutions discriminating different eukaryotic groups (Figure 1 C). Nearly all metazoa (93 %, including 98 % of mussels), fungi (95 %) and cercozoans (80 %) showed a TAA, while nearly all remaining eukaryotic groups were distinguished by GAG or GTG at the according position. We thus designed three blocking primers (DreissDiet_TAA_R, DreissDiet_GAG_R, DreissDiet_GTG_R), which were distinguished at the 4th-6th nucleotide position and should discriminate between these different eukaryotic groups.

However, these primers will not generally block amplification of other metazoan filter feeders. To assess the diet of a broader spectrum of filter feeding animals, we used the three discriminating nucleotide positions at the 4th-6th position of the *Dreissena* primer. We designed three additional primers, which started at these positions. One of these primers (Metazoa_TAA_R: 3’-TAA) should efficiently amplify metazoans, fungi and cercozoans, while blocking all other taxa. The primer will result in up to three 3’-mismatches (A-G, T-A & T-G), which will strongly affect amplification efficiency. The other two (NonMetazoaDiet_GAG_R: 3’-GAG or NonMetazoaDiet_GTG_R: 3’-GTG) will amplify the majority of other eukaryotes, while reducing amplification of metazoans, fungi and cercozoans due to up to three 3’-mismatches (T-C, A-A & A-C). These latter two primers should thus be well suited to enrich non-metazoan eukaryotic taxa from extracts of filter feeding metazoans. In comparison to the *Dreissena* primers, however, the blocking should be less efficient, as they only show 2-3 3’-mismatches.

Corresponding forward primers were designed at the 3’-end of the V8 region (Figure 1a). The first forward primer (Euk_F1) was universal for eukaryotes and was designed as a complementary primer to the metazoan primers (DreissDiet_TAA_R, Metazoa_TAA_R). The second primer (NonMetazoa_F1) discriminated bivalves and other metazoans from other eukaryotes by its first base (Figure 1 a). While all bivalves (100 %) and the majority of metazoans (78.9 %) showed a G at that position, nearly all other eukaryotes showed a T (99-100 % depending on group), resulting in a T-C mismatch with bivalves. This primer was used as complement to the non-metazoan reverse primers (DreissDiet_GAG_R, DreissDiet_GTG_R, NonMetazoaDiet_GAG_R & NonMetazoaDiet_GTG_R). The resulting PCR fragments of these primer combinations covered the complete V8 and V9 region and reached a length of 360 bp on average. DNA in predators’ guts and eDNA is often degraded and present in small fragment sizes (Krehenwinkel et al. 2017). The relatively long fragment length amplified by our primer combinations may therefore make them unsuitable for some of this highly degraded DNA. To enable the recovery of degraded DNA fractions, we designed an additional eukaryote specific forward primer (Euk_F2) in the conserved DNA stretch between V8 and V9. This primer can be combined with all other reverse primers and amplifies a fragment length of 161 bp on average. For an overview of all designed primer combinations and their blocking efficiency see Table 1).

All primer pairs were tested with different eukaryotic taxa in gradient PCRs at annealing temperatures from 50-60 °C, to estimate the optimal annealing temperature. PCRs were run in 10 µl volumes and with 35 cycles using the Qiagen Multiplex Kit according to the manufacturer’s protocols (Qiagen, Hilden, Germany). PCR success was checked on a 1.5 % agarose gel.

### Test of the designed PCR primers

Out of the 9 total primers we designed, 7 were relevant to test their efficiency in blocking bivalve amplification. These were further tested in the following experiments. We first tested the efficiency of the *Dreissena* blocking primers in recovering a broad taxonomic diversity of eukaryotes, while at the same time suppressing *Dreissena* amplification. For this purpose, we collected samples of zebra and quagga mussels from two sites in the river Danube. In addition, we collected samples of the zebra mussel from the lake Stechlin. The mussels were collected from hard substrate close to the shore by hand and subsequently stored in the gas phase over liquid nitrogen. Zebra and quagga mussels were identified morphologically using a standardized protocol (Teubner et al. 2016) and stored separately. Before the collection of the mussels, two water samples of 1 liter each were collected at the same location where the mussels were collected. These samples served as a direct comparison of the eDNA present in the water column and the recovered DNA population from mussels. The water samples were filtered using a nitrocellulose filter of 0.45 μm pore size (Thermo Fisher Scientific Inc., Waltham, USA). DNA was then extracted from the filter using the Qiagen DNeasy PowerWater Kit according to the manufacturer’s protocol.

The mussels were dissected using the standardized protocol of the ESB (Teubner et al. 2018). Each mussel was briefly thawed and opened with tweezers, and the tissue and water were carefully removed. We then finely ground the entire mussel in a mortar while adding liquid nitrogen. To maximize taxon recovery, we combined the tissue of 32 mussels of each species for the two collection sites at the Danube. The two mussel species were processed separately. To estimate the recovery of prey diversity with fewer mussels, we also included an extraction of 4 mussels from the lake Stechlin. To avoid cross contamination, the mortar was thoroughly cleaned and treated with bleach after each processed sample.

The homogenate was transferred to a tube and DNA extracted from it using the Qiagen Puregene DNA extraction kit according to the manufacturer’s protocols. We did not isolate DNA from the complete lysate volume, instead we used 450μl, as even in small subvolumes of larger lysates the DNA represent community composition well (Creedy et al. 2019).

The water eDNA samples and the mussel samples were then amplified with different *Dreissena* blocking primer pairs (EukF1 + DreissDiet_TAA_R, NonMetazoaF1 + DreissDiet_GAG_R, NonMetazoaF1 + DreissDiet_GTG_R), each using the Qiagen Multiplex PCR kit as described above at an optimal annealing temperature of 55 °C. After the first PCR, a second indexing PCR of 5 cycles was added, as described in Krehenwinkel et al. (2018) and using the same reaction conditions as in the first round PCR. The second round PCR products were visualized on a 1.5 % agarose gel, pooled in approximately equal concentrations based on gel band intensity and then cleaned of leftover primer with 1 X AmPure Beads XP (Beckman & Coulter, California, USA).

To test the general metazoan blocking primers, which allow the enrichment of non-metazoan prey from filter feeding animals (NonMetazoaF1 + NonMetazoaDiet_GAG_R, NonMetazoaF1 + NonMetazoaDiet_GTG_R), we tested a different mussel species, the blue mussel *Mytilus edulis*, which does not show the characteristic 3’-TC mismatch of *Dreissena* spp.. We used a selection of three blue mussel samples from each of two sites: the Northern Sea and Baltic Sea of Germany. Each sample consisted of several 100 mussels collected throughout the year. The mussels were sorted, measured, dissected and then homogenized and stored on liquid nitrogen (Paulus et al. 2018). DNA was extracted from 50 mg of homogenate using the Qiagen Puregene Kit, and the samples were then amplified using the two primer pairs as described above. Negative control PCRs were run alongside all samples.

### Application of our protocol using a Dreissena polymorpha time series of ESB homogenates

To show a direct application of our protocol, we analyzed time series samples of the zebra mussel, collected by the ESB in three German rivers following the guideline of Teubner et al. (2018). We included samples from the rivers Rhine, Elbe and Saar, which were collected between 1994 and 2016. We extracted DNA from ∼50 mg of homogenate of each sample, using the Qiagen PureGene extraction kit as described above. All samples were amplified for two *Dreissena* blocking primer pairs (EukF1 + DreissDiet_TAA_R, NonMetazoaF1 + DreissDiet_GTG_R) and libraries prepared and pooled as described above. Negative control PCRs were run alongside all samples.

### Sequencing and sequence analysis

The two resulting libraries were quantified using a Qubit Fluorometer (Thermo Fisher Scientific Inc., Waltham, USA), pooled in equimolar proportions and then sequenced on an Illumina MiSeq using V3 chemistry and 600 cycles (Illumina Inc., San Diego, USA). The samples were demultiplexed using CASAVA v1.8.2 (Illumina Inc.) with no mismatches allowed. Demultiplexed fastq files were merged using PEAR (Zhang et al. 2014) with a minimum overlap of 50 and a minimum quality of 20. The merged reads were additionally filtered for a minimum quality of Q30 over > 90 % of the sequence and then transformed to fasta files using the FASTX-Toolkit (Gordon & Hannon 2010). Primer sequences were then trimmed off using *sed* in UNIX scripts. The amplicons resulting from our different primers were largely overlapping, but showed slight differences in length (see Figure 1). We thus also trimmed all amplicons to the same starting and end position using *sed*. The processed reads were then dereplicated and clustered into zero radius OTUs (hereafter OTUs) using USEARCH (Edgar 2010). A minimum cluster size of 8 was used. A de novo chimera removal is included in the clustering pipeline. As the 18S gene is rather conserved, we refrained from clustering into 3 % radius OTUs, as this would have likely deflated true taxonomic diversity. The recovered OTU richness for 18SrDNA zero radius OTUs corresponds relatively well with that found with 3 % radius OTUs used in mitochondrial COI for metazoans (de Kerdrel et al. 2020). All resulting OTUs were searched against the complete NCBI nucleotide database (downloaded 02/2020) using BLASTn (Altschul et al. 1990). Taxonomy was then assigned to the resulting BLAST output using a custom *python* script (de Kerdrel et al. 2020). An OTU table was then constructed for all samples using USEARCH. For our comparative analysis of *Dreissena* and water eDNA samples, we rarefied the OTU table for all samples to 10,000 reads per sample, at which saturation of diversity was reached (Suppl. Figure 1).

We first tested the efficiency of our primers for suppressing amplification of *Dreissena* mussels. For this purpose, we calculated the proportion of recovered non-mussel sequences for each sample based on the previous taxonomic assignments. We also explored the OTU table for possible parasites and commensals of the mussels, which should also not be treated as diet, and estimated their proportion among the total reads. The OTU table was then used to calculate alpha and beta diversity within and between samples using VEGAN (Oksanen et al. 2013) in R v4.03 (R Core Team 2013) & RStudio v1.4.1103 (RStudio Team 2020). Factors influencing beta diversity were estimated using a PERMANOVA using the *adonis* function in *vegan*. Community similarity was visualized using NMDS plots based on the metaMDS function in vegan (distance = “jaccard”, binary = T, k = 3, trymax = 999). All plots in this study were created with GGPLOT2 (Wickham 2016). We also compared the recovered taxonomic composition and diversity in mussel and filtered water eDNA samples and between the different primer pairs used in our experiments. Using the time series samples, we analyzed changes in the mussels’ diet composition within rivers over time and between rivers.

## Results

### Blocking performance of the designed PCR primers

We recovered an average of 45,376 reads for each of our *Dreissena* and water samples and 7,605 reads for the *Mytilus* samples. After quality filtering and OTU clustering, we found a total of 6012 OTUs in the dataset.

Our primers performed quite differently in the recovery of non-mussel sequences. The *Dreissena* primers were all highly efficient at blocking out the mussels’ sequences. For the primers EukF1 + DreissDiet_TAA_R, on average 82.11%, 99.9% for NonMetazoaF1 + DreissDiet_GAG_R and 99.6% for NonMetazoaF1 + DreissDiet_GTG_R of the sequences recovered were non-mussel sequences (Figure 2a). The recovered OTU richness was comparable between the primers. Mussel samples recovered 199 OTUs on average for primers EukF1 + DreissDiet_TAA_R, 206 for NonMetazoaF1 + DreissDiet_GAG_R and 153 for NonMetazoaF1 + DreissDiet_GTG_R. A combination of the three primers into a single sample led to a considerable increase to 343 recovered OTUs. As expected, the primer pairs that did not contain the additional 3’-TC blocking sequence (NonMetazoaF1 + NonMetazoaDiet_GAG_R, NonMetazoaF1 + NonMetazoaDiet_GTG_R) amplified considerably more mussel sequences. An average of 25.35 % and 33.99 %, respectively, of the total reads were identified as non-blue mussel sequences for these primers (Suppl. Figure 2).

**Figure 2).**
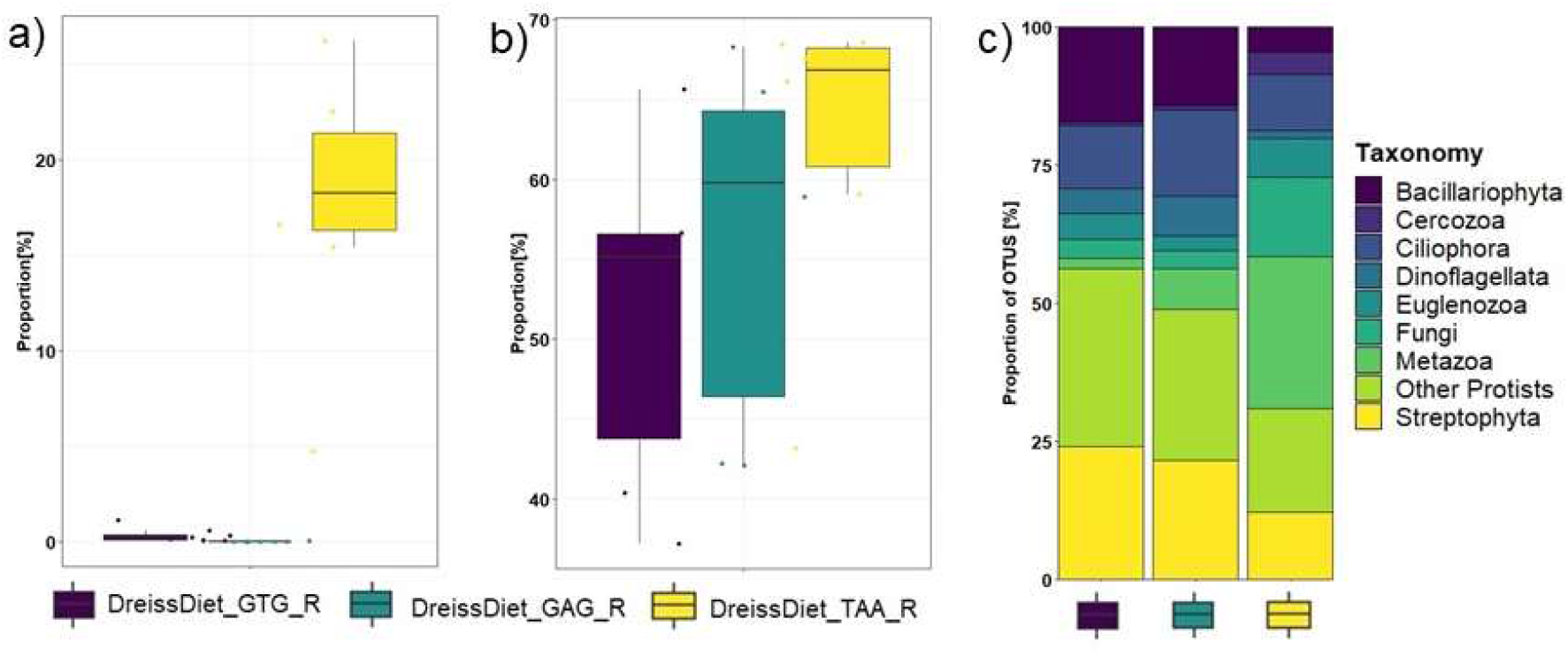
a) Recovered proportion of Dreissena reads for three Dreissena blocking primer combinations b) Recovered proportion of Dreissena reads and parasite or commensal reads for the same three primer combinations c) Higher taxonomic composition of the OTUs recovered by the three primer pairs.

We also found a total of 106 OTUs of likely commensals and parasites of the mussels in the data, for example the ciliate *Conchopipthurus* sp. and several trematodes like *Cryptocoyle lingua* and *Aspidogaster conchicola*. As such taxa are often quite abundant in their host, they made up a considerable proportion of the reads. After removing these taxa from the read population, 62.17 % of potential dietary reads remained for the EukF1 + DreissDiet_TAA_R primers, 56.29 % for NonMetazoaF1 + DreissDiet_GAG_R and 51.69 % for NonMetazoaF1 + NonMetazoaDiet_GTG_R, on average (Figure 2b).

The recovered taxonomic composition of OTUs was comparable between the NonMetazoaF1 + DreissDiet_GAG_R and NonMetazoaF1 + DreissDiet_GTG_R primers. A clear difference in the taxonomic composition of recovered OTUs was found between the previously mentioned primers and the EukF1 + DreissDiet_TAA_R primers. While the first two recovered considerably more plants (mostly green algae) and Bacillariophyta, the latter showed more Metazoa and Cercozoa OTUs (Figure 2c).

### Comparison of recovered communities by mussels and eDNA water samples

The three *Dreissena* diet primer pairs were merged into a single dataset for each sample for this analysis and the 106 mussel parasite OTUs removed from the data. Mussel and water samples recovered comparable compositions of higher taxa in our analysis (Figure 3a). The only noteworthy difference between water and mussel samples was found for the recovery of metazoa in Kelheim and Jochenstein, which were more taxonomically rich in mussels than water samples (18.11 ± 6.635 % vs. 5.93 ± 1.205%). The comparable taxonomic composition recovered by water and mussel samples held true when lower taxonomic ranks were compared. For example, very similar proportions of OTUs were recovered from different metazoan phyla between water and mussel samples (Figure 3b). A slight difference was found for nematodes, which showed more OTUs in mussel samples. Moreover, we found a larger proportion of rotifer OTUs in water samples from Kelheim than in mussel samples.

**Figure 3.**
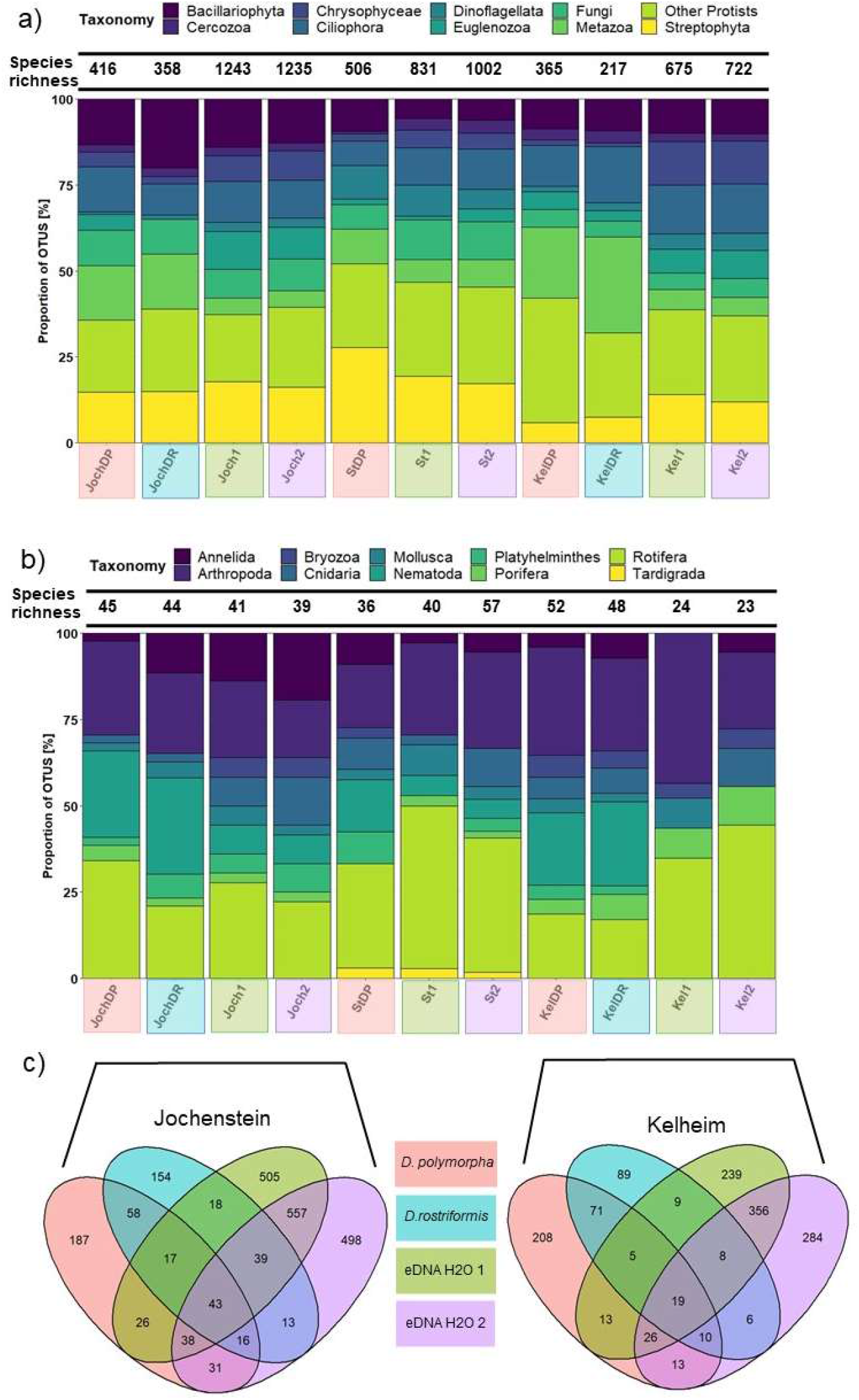
a) Higher taxonomic composition of the recovered OTUs comparing water eDNA samples, Dreissena polymorpha (DP) and D. rostriformis (DR) DNA extracts from the different sites (Jochenstein = Joch, Kelheim = Kel, Stechlin = St). The plots show the total number of recovered OTUs in each sample on top and proportion of different taxa among these OTUs. b) Plot showing phylum level taxonomic composition of the recovered metazoan OTUs for the same samples and sites. c) Venn diagrams showing the recovered OTU numbers and their overlap between D. polymorpha, D. rostriformis and the two eDNA water samples for the two sites at the Danube river.

Across all eukaryotes, the water samples recovered a significantly higher OTU richness than the mussel samples (953 vs. 343 on average, t-test, *P* < 0.05) (Figure 3 c + d & Suppl. Figure 3). The two mussel species *D. polymorpha* and *D. rostriformis* recovered relatively similar richness (281 vs. 345 on average). Comparing the two sites Kelheim and Jochenstein, mussel and water samples showed similar trends of recovered richness, with both showing higher richness in Kelheim than in Jochenstein. The number of mussels in a sample did not appear to significantly contribute to the number of recovered species. The samples of *D polymorpha* from Lake Stechlin (only 4 mussels) recovered an even higher OTU richness than the other two samples of 32 mussels (506 vs 391). Interestingly, mussel samples appeared as efficient as water samples in recovering metazoans. On average, mussels recovered 45 metazoan OTUs, water samples 37. The difference was not significant (t-test, P > 0.05).

While they recovered relatively similar communities at higher taxonomic levels, we found quite distinct communities at the OTU level for mussel and water samples (Figure 3 c + d & Suppl. Figure 3 & 5). Only 28.25 % of the OTUs found in *D. polymorpha* were also found in water samples, likewise 33.53 % of the OTUs in *D. rostriformis*. But even different water samples and the two analyzed mussel species at each site recovered different OTU compositions. On average, 57.53 % of the recovered OTUs were shared between two water samples and 34.01 % between *D. polymorpha* and *D. rostriformis* at the same site.

The relatively small overlap of recovered OTUs between water and mussel samples was well reflected in our NMDS plot (Figure 4). Here, water and mussel samples were well separated on the first axis. However, the NMDS also distinguished the different sampling sites. The general pattern of differentiation was comparable between water samples and mussels. Samples from Lake Stechlin formed a separate cluster, distant from the two samples from the Danube river. The two samples from Jochenstein and Kelheim (both Danube river) were also separated by water and mussel eDNA, grouping together closely. The separation of Kelheim and Jochenstein was less clear for the mussels, than for the water samples (Figure 4a). However, when mussel parasite OTUs were included in the dataset, the pattern reversed, with the two sites from the Danube being more distinct in mussels (Figure 4b). An inclusion of mussel parasites did not have any effect on the differentiation of the water samples. The mussel eDNA samples also showed a differentiation between the recovered communities of *D. rostriformis* and *D. polymorpha*. The different taxonomic compositions recovered for the two mussel species were supported by our NMDS plot, particularly, when mussel parasites were included in the data (Suppl. Figure 4). The recovered differentiation in the NMDS plot was also supported by a PERMANOVA, suggesting significant (P < 0.001) effects of sample type (R2 = 0.193), sampling site (R2 = 0.199), mussel species (R2 = 0.119) on the recovered beta diversity pattern. No significant difference was found for the communities recovered by the three different primer sets (R2 = 0.053, P > 0.05)

**Figure 4.**
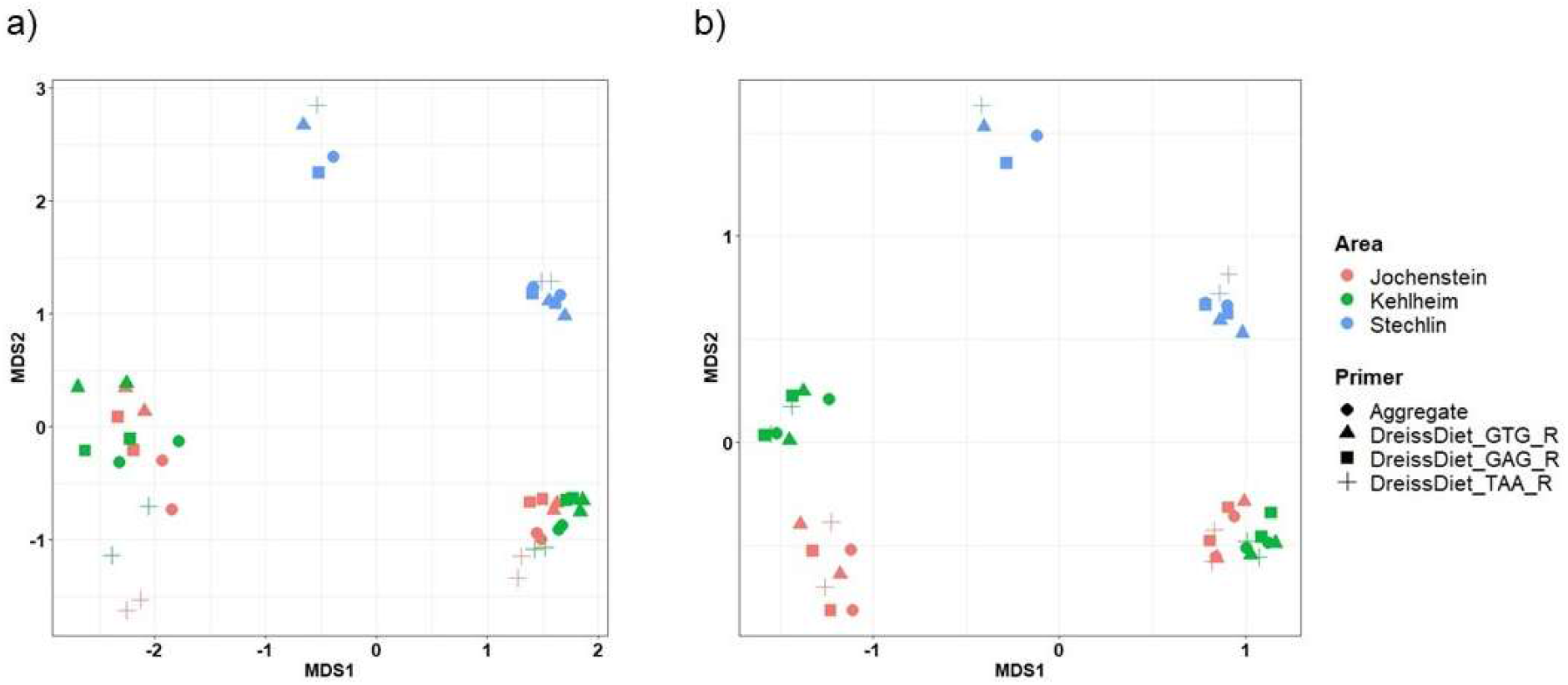
a) NMDS plots based on Jaccard dissimilarity showing community differentiation of eDNA water samples and Dreissena mussel samples for the two sites ate the Danube (Kelheim and Jochenstein) and Lake Stechlin. The first axis separates water (right) and mussels (left). The shapes distinguish the different primer combinations used, as well as a combination of all three into a merged dataset (Aggregate). a) Shows the NMDS plot (stress = 0.088), with commensal and parasite OTUs of the mussels removed, b) (stress = 0.089) shows the same plot including parasites and commensals. For a distinction between D. polymorpha and D. rostriformis associated communities, see Suppl. Figure 4a & b.

### Application of our protocol using a Dreissena polymorpha time series ESB homogenates

The ESB homogenate samples recovered 28,509 reads on average and a total of 1716 OTUs. Due to their lower coverage, we rarefied these samples to 5,000 reads each and then merged the dataset for EukF1 + DreissDiet_TAA_R, and NonMetazoaF1 + DreissDiet_GTG_R. Hence, the final dataset also had a coverage of 10,000 reads per sample. We did not recover significant differences in richness between the three sampled rivers (ANOVA, *P* > 0.05). On average, we found 121 OTUs in the samples from the Rhine, 138 in those from the Saar and 107 in those from the Elbe. For Rhine and Saar, we also did not find changes of richness between different sampling periods. For the Elbe, however, we found a significant decline of richness with time from 140 OTUs in 1998 to only 63 in 2016 (Linear model, R^2^ = 0.89, *P* < 0.05).

The different rivers were well separated by our NMDS plot (Figure 5a). A PERMANOVA suggests that sampling site primarily contribute to the observed beta diversity (R^2^ = 0.22, *P* < 0.001). However, we also found a strong positive correlation between beta diversity between samples and the temporal distance between sampling events (Linear model, *P* < 0.05, R^2^_Rhine_ = 0.73, R^2^_Elbe_ = 0.85) for the river Rhine and Elbe (Figure 5b). The community composition in these rivers strongly turned over in the past 20 years. This effect was not visible for the Saar river.

**Figure 5.**
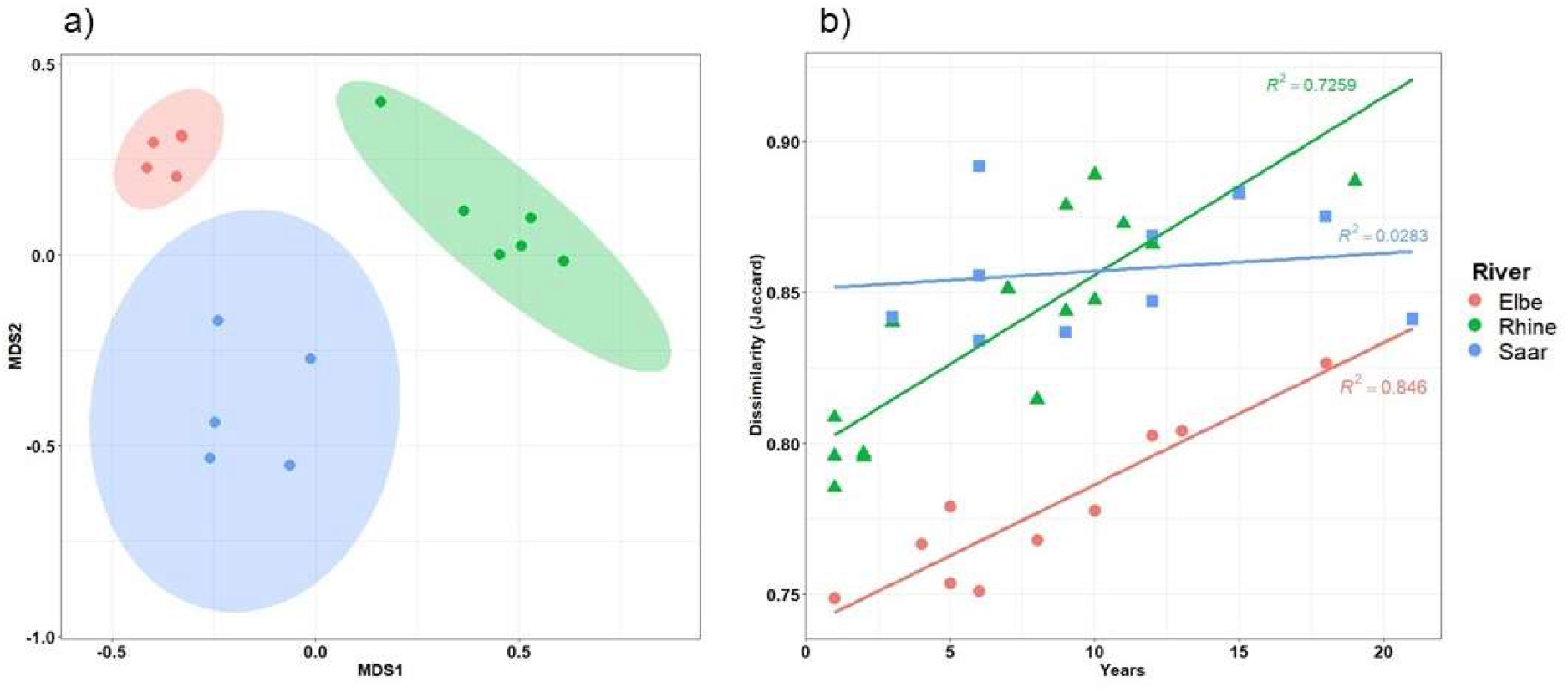
a) NMDS (stress =0.080) plot based on Jaccard dissimilarity showing differentiation of Dreissena polymorpha associated eukaryotic communities between ESB homogenates collected from three different rivers. Ellipses represent the 95% confidence interval. b) Correlation of Jaccard dissimilarity between the D. polymorpha associated communities and the number of years between sampling events.

## Discussion

### A primer set for high throughput diet analysis and parasite monitoring in mussels

In the past years, metabarcoding of gut content has developed into an increasingly popular tool for food web analysis (Kennedy et al. 2020). However, this method often comes with a tradeoff between maximizing the recovered prey diversity and minimizing the amplification of predator DNA (Krehenwinkel et al. 2017). Here we show that the use of primers with broad taxonomic specificity, which incorporate 3’-end nucleotide mismatches between prey and predator, is an efficient way to tackle this issue (Stadhouders et al. 2010; Krehenwinkel et al. 2019). The primer set we introduce here allows for a near complete suppression of the amplification of mussels of the genus *Dreissena* and 33 other bivalve genera in 16 families. At the same time, our assay recovers a very broad spectrum of potential eukaryote dietary taxa. Based on additional group specific mismatches, we designed three reverse primers to each target slightly different eukaryotic groups. These primers amplify nearly identical DNA fragments and could thus also be used in a multiplex reaction (Krehenwinkel et al. 2019) to reduce the necessary number of PCRs. Our protocol opens up new possibilities to study aquatic food webs, in which filter feeding mussels often play a critical role (Newell 2004; Vaughn et al. 2008). Moreover, the marker set will be of great value for studying the impact of invasive *Dreissena* mussels on aquatic ecosystems (Maguire & Grey 2006; Miller & Watzin 2007).

It is noteworthy that not all taxa recovered from the homogenized mussels are likely to be their actual diet. Mussels filter the water column for most particles in a certain size range (Sprung & Rose 1988). A smaller part of the recovered DNA molecules would probably have not been consumed, but rejected as pseudofeces. However, considering that our analysis includes the complete digestive tract of the mussel, the larger proportion of recovered taxa is probably the mussel’s actual diet. In the case of zooplankton feeding mussels, another source of bias could be secondary predation (Sheppard et al. 2005), e.g., DNA from phytoplankton ingested by the mussel’s zooplankton prey. And last, we identified several parasitic and commensal organisms living in close association with the mussels (Molloy et al. 1997). Such taxa should be removed before actual diet analysis is performed. At the same time, the possibility to monitor parasites with our method offers an interesting application for monitoring health in natural mussel populations or in mussel aquaculture, which is of great economic importance in some areas (Shumway et al. 2003). In our analysis, these parasites and commensals were barely detected in water eDNA samples, collected in close proximity to the mussels. For the monitoring of pathogens in mussel populations, the collection of mussel tissue is thus inevitable. Another interesting finding was the more pronounced differentiation of sampling sites in the Danube river, when parasite OTUs were included. This suggests that *Dreissena* mussels in Germany may harbor site specific parasite communities at a very small geographic scale.

An interesting result from our analysis in sympatric *D. polymorpha* and *D. rostriformis* is their possible selective feeding. While we found some overlap in the consumed taxa, a considerable proportion of dietary organisms were not shared. This is surprising considering that the species were sampled next to each other along river embankments. The reason for the selectivity could be the slightly different size of the mussel species and associated particle size of the filtered dietary items. But it could also be based on other factors inherent to the two species. It has long been assumed that the quagga mussel is replacing the zebra mussel in its current invasie range (Mathews et al. 2014). However, a niche partitioning by particle selectivity and following different dietary resource use (Baker & Levinton 2003) could allow a long-term coexistence of the two species. A particularly strong difference was also found for parasite communities between *D. polymorpha* and *D. rostriformis*, suggesting that site and species specific parasites may colonize invasive *Dreissena* populations.

It should be noted that for small mussels like *Dreissena* spp., bacteria may also be an important part of the diet (Dionisio Pires 2005). We did not analyze this here, as we were particularly interested in eukaryotic taxa. Bacterial composition in mussels could be easily scored using 16SrDNA primers and corresponding molecular protocols (Kozich et al. 2013).

While our marker set has a broad amplification success across bivalves, it blocks very few other filter feeding metazoans. The additional primers we tested in *Mytilus edulis* show fewer mismatches with this bivalve. It thus is less efficient in blocking the mussel’s DNA from amplification. However, with constantly dropping sequencing costs, even a 30 % recovery of possible dietary taxa is a reasonable recovery rate. As these primers block all metazoans, dietary analysis would have to be limited to the non-metazoan parts of the planktonic community. But at the same time, this makes the primers well suited to score non-metazoan dietary components of any filter feeding aquatic animals, e.g. sponges, cnidarians, annelids or fish, which will all be suppressed from amplification.

### The utility of mussels as biological eDNA filters

The filter feeding mechanism of many aquatic metazoans is quite similar to the commonly used filtering of water for eDNA analysis (Barnes et al. 2020; Wilcox et al. 2015; Jo et al. 2019). This has led to the suggestion that aquatic filter feeders could be used as biological eDNA collectors. Recent work in sponges (Mariani et al. 2019) suggests that they can indeed serve this purpose, for example in fish monitoring (Turon et al. 2020). Even gut content analysis of detritus feeders like shrimps has made it possible to reconstruct fish communities (Siegenthaler et al. 2019). While these results are promising, the utility of these organisms as eDNA filters and the taxonomic breadth of recovered taxa are limited. For example, the above studies were mostly limited to the detection of fish. Such taxa, which are quite evolutionarily distant from the consumer, can be enriched from the DNA extract with taxon-specific primers. However, a broad-spectrum analysis of aquatic biodiversity is not possible due to the overabundance of the consumer’s DNA (Krehenwinkel et al. 2017). Our protocol addresses this problem by providing a set of primers that explicitly blocks the amplification of consumer DNA while retaining the ability to amplify a broad taxonomic range of eukaryotes.

Mussels should be well suited as biological eDNA filters. Their efficient filtering of plankton (Fanslow et al. 1995) is dramatically illustrated by the strong cascading effect of the invasion of *Dreissena* mussels in American lakes on limnic ecosystems (Lavrentyev et al. 1995; MacIsaac et al. 1995; Maguire & Grey 2006). Their efficacy of particle removal has even led to the suggestion to use them as biofilters for water clarification (Elliot et al. 2008).

Our results show that mussels retain a high diversity of aquatic eukaryotes as potential dietary items. The recovered taxa also show a considerable overlap with water samples collected next to the mussel colonies. The alpha and beta diversity trends recovered between different sites by water samples were also reflected in the mussel data. However, the recovered richness was considerably higher in the water samples, suggesting a certain selectivity of the mussel’s filtering apparatus (Baker & Levinton 2003). Only a subset of particles retained by a nitrocellulose filter were detectable in mussels. This could be the consequence of the filter’s ability to retain finer particle sizes. Recent work suggests that eDNA particles are present in diverse size ranges, spanning from < 0.2 toseveral 100 µm, but are particularly abundant at the lower end of the spectrum (Turner et al. 2014). Different studies recommend different filter size from 10 µm down to 0.2 µm (Barnes et al. 2020; Wilcox et al. 2015; Jo et al. 2019). Here we used a filter pore size of 0.45 µm. *Dreissena* mussels show an imperfect retention of particles larger than 0.7 µm and a total retention only at particle sizes larger than 5 µm (Sprung & Rose 1988). The retention efficiency shifts to larger particles in larger mussels, like the blue mussels (*Mytilus* spp.), where only particles > 7 µm show a total retention (Strohmeier et al. 2012). Our mussel-based analysis may thus have omitted some small particles. As the retention of mussels correlates with their size (MacIsaac et al. 1995), a broader spectrum of dietary taxa could possibly be recovered by choosing mussels of various sizes for the eDNA analysis, including very small specimens.

On the other hand, a noteworthy proportion of taxa which we detected in mussels was not found in water samples and mussel preformed equally well in recovering metazoans. Many of these may be taxa that live in close association with the mussels, e.g. parasites and commensals (Molloy et al. 1997). However, some of these may also be species that do not release much eDNA into the water column. It is well known that water based eDNA analysis does not necessarily yield an exhaustive overview of the local community (Koziol et al. 2019). It is also quite likely that the mussels can recover taxa over a broader timescale. While non-palatable items are expelled as pseudofeces, the ingested dietary particles will pass through the gut for several hours before being expelled (Hawkins et al. 1990). DNA molecules can therefore be recovered even during the digestion process (Krehenwinkel et al. 2019). The recovered dietary DNA spectrum thus likely reflects the result of up to a day of filtering. A filtered water sample, in contrast, is just a snapshot of the present eDNA at a certain site.

An ideal eDNA sample should be taxonomically exhaustive, while at the same time minimizing processing effort. A water sample represents an ideal compromise between these two demands. Yet, the use of DNA extracts from filter feeders adds another complementary perspective to eDNA analysis. A relatively small number of mussels is probably sufficient to serve as an eDNA sample, and their processing in the laboratory is quite straightforward. Nevertheless, the ease of handling and processing of water samples makes it unlikely that they will be replaced by alternative eDNA strategies. However, filter feeding organisms could be of great utility in some instances. For example, in biological surveys of sea floors, mussels are a common bycatch. These specimens could be immediately used as promising eDNA sources for deep sea habitats, without the need for taking additional water samples.

### An application of our protocol to time series samples

Our analysis of time series samples shows an interesting application of our protocol. As for the previous analysis on freshly collected mussels, the time series samples recovered different taxa in different sampling sites, most likely as a result of differences in plankton community compositions in the different rivers. Moreover, most sampling sites suggest a pronounced temporal turnover of the recovered dietary community. In the past decades, decreasing eutrophication and increasing warming have likely led to state shifts in rivers, which strongly affect the plankton communities (Abonyi et al. 2018). Another possible effect on the recovered dietary community could be the taxonomic composition of the analyzed mussel samples. While ESB samples from before 2004 are entirely composed of *D. polymorpha*, some samples may contain small numbers of *D. rostriformis*, after this species began its invasion into German waterways around 2004 (Paulus et al. 2014). The two species may show different selectivity for dietary particles, which could also explain the observed turnover. To disentangle these effects, we are currently working on a more detailed analysis of the *Dreissena* time series.

## Acknowledgements

We thank Karin Fischer for help with laboratory work and Bernhard Fontaine, Gleb Stalsky and Alois Deutsch for providing support during processing of samples. Niklas Richter prepared the ESB DNA extractions we used here. We thank the German Environment Agency and the German Environmental Specimen Bank for providing the time series samples of *Dreissena polymorpha*.

## Author contributions

HK devised the study, DT, MW and LB performed the field work. SW and LB performed laboratory work. HK and SW analyzed the data and wrote the manuscript. All authors have read and approved the final version of the manuscript.

## Data accessibility statement

All alignments of 18SrDNA sequences for primer design and of the priming sites for different eukaryotic taxa, all raw reads and OTU tables will be made available on the Dryad Digital Repository after acceptance of the manuscript.

## References

Abonyi, A., Ács É., Hidas A., Grigorszky, I., Várbíró, G., Borics, G., & Kiss, K. T. (2018). Functional diversity of phytoplankton highlights long-term gradual regime shift in the middle section of the Danube River due to global warming, human impacts and oligotrophication. Freshwater Biology, 63(5), 456–472. https://doi.org/10.1111/fwb.13084

Albaina, A., Aguirre, M., Abad, D., Santos, M., & Estonba, A. (2016). 18S rRNA V9 metabarcoding for diet characterization: A critical evaluation with two sympatric zooplanktivorous fish species. Ecology and Evolution, 6(6), 1809–1824. https://doi.org/10.1002/ece3.1986

Altschul, S. F., Gish, W., Miller, W., Myers, E. W., & Lipman, D. J. (1990). Basic local alignment search tool. Journal of Molecular Biology, 215(3), 403–410.

Baker, S. M., & Levinton, J. S. (2003). Selective feeding by three native North American freshwater mussels implies food competition with zebra mussels. Hydrobiologia, 505(1), 97–105. https://doi.org/10.1023/B:HYDR.0000007298.52250.99

Barnes, M. A., Turner, C. R., Jerde, C. L., Renshaw, M. A., Chadderton, W. L., & Lodge, D. M. (2014). Environmental conditions influence eDNA persistence in aquatic systems. Environmental Science and Technology, 48(3), 1819–1827. https://doi.org/10.1021/es404734p

Burrows, M. J. (2013). Invading dreissenid mussels transform the 100-year-old international joint commission. Quagga and Zebra Mussels: Biology, Impacts, and Control, Second Edition, 8(1), 187–194. https://doi.org/10.1201/b15437

Capra, E., Giannico, R., Montagna, M., Turri, F., Cremonesi, P., Strozzi, F., … Pizzi, F. (2016). A new primer set for DNA metabarcoding of soil Metazoa. European Journal of Soil Biology, 77, 53–59. https://doi.org/10.1016/j.ejsobi.2016.10.005

Choi, J., & Park, J. S. (2020). Comparative analyses of the V4 and V9 regions of 18S rDNA for the extant eukaryotic community using the Illumina platform. Scientific Reports, 10(1), 6519. https://doi.org/10.1038/s41598-020-63561-z

Connelly, N. A., O’Neill, C. R., Knuth, B. A., & Brown, T. L. (2007). Economic impacts of zebra mussels on drinking water treatment and electric power generation facilities. Environmental Management, 40(1), 105–112. https://doi.org/10.1007/s00267-006-0296-5

Creedy, T. J., Ng, W. S., & Vogler, A. P. (2019). Toward accurate species-level metabarcoding of arthropod communities from the tropical forest canopy. Ecology and Evolution, 9(6), 3105–3116. https://doi.org/10.1002/ece3.4839

de Kerdrel, G. A., Andersen, J. C., Kennedy, S. R., Gillespie, R., & Krehenwinkel, H. (2020). Rapid and cost-effective generation of single specimen multilocus barcoding data from whole arthropod communities by multiple levels of multiplexing. Scientific Reports, 10(1), 1–12. https://doi.org/10.1038/s41598-019-54927-z

Dionisio Pires, L. M., Bontes, B. M., Van Donk, E., & Ibelings, B. W. (2005). Grazing on colonial and filamentous, toxic and non-toxic cyanobacteria by the zebra mussel Dreissena polymorpha. Journal of Plankton Research, 27(4), 331–339. https://doi.org/10.1093/plankt/fbi008

Edgar, R. (2010). Usearch. Lawrence Berkeley National Lab.(LBNL), Berkeley, CA (United States).

Elbrecht, V., & Leese, F. (2017). Validation and development of COI metabarcoding primers for freshwater macroinvertebrate bioassessment. Frontiers in Environmental Science, 5(APR), 11. https://doi.org/10.3389/fenvs.2017.00011

Elliott, P., Aldridge, D. C., & Moggridge, G. D. (2008). Zebra mussel filtration and its potential uses in industrial water treatment. Water Research, 42(6–7), 1664–1674. https://doi.org/10.1016/j.watres.2007.10.020

Fanslow, D. L., Nalepa, T. F., & Lang, G. A. (1995). Filtration Rates of the Zebra Mussel (Dreissena polymorpha) on Natural Seston from Saginaw Bay, Lake Huron. Journal of Great Lakes Research, 21(4), 489–500. https://doi.org/10.1016/S0380-1330(95)71061-9

Fernández, A., Grienke, U., Soler-Vila, A., Guihéneuf, F., Stengel, D. B., & Tasdemir, D. (2015). Seasonal and geographical variations in the biochemical composition of the blue mussel (Mytilus edulis L.) from Ireland. Food Chemistry, 177, 43–52. https://doi.org/10.1016/j.foodchem.2014.12.062

Giebner, H., Langen, K., Bourlat, S. J., Kukowka, S., Mayer, C., Astrin, J. J., … Fonseca, V. G. (2020). Comparing diversity levels in environmental samples: DNA sequence capture and metabarcoding approaches using 18S and COI genes. Molecular Ecology Resources, 20(5), 1333–1345. https://doi.org/10.1111/1755-0998.13201

Gordon, A., & Hannon, G. J. (2010). FASTX-TOOLKIT, version 0.0. 14. Computer Program and Documentation Distributed by the Author

Gregorič, M., Kutnjak, D., Bačnik, K., Gostinčar, C., Pecman, A., Ravnikar, M., & Kuntner, M. (2020). Spider webs as eDNA tool for biodiversity assessment of life’s domains. BioRxiv. https://doi.org/10.1101/2020.07.18.209999

Hawkins, A. J. S., Navarro, E., & Iglesias, J. I. P. (1990). Comparative allometries of gut-passage time, gut content and metabolic faecal loss in Mytilus edulis and Cerastoderma edule. Marine Biology, 105(2), 197–204. https://doi.org/10.1007/BF01344287

Hoogendoorn, M., & Heimpel, G. E. (2001). PCR-based gut content analysis of insect predators: Using ribosomal ITS-1 fragments from prey to estimate predation frequency. Molecular Ecology, 10(8), 2059–2067. https://doi.org/10.1046/j.1365-294X.2001.01316.x

Jo, T., Murakami, H., Yamamoto, S., Masuda, R., & Minamoto, T. (2019). Effect of water temperature and fish biomass on environmental DNA shedding, degradation, and size distribution. Ecology and Evolution, 9(3), 1135–1146. https://doi.org/10.1002/ece3.4802

Kennedy, S. R., Prost, S., Overcast, I., Rominger, A. J., Gillespie, R. G., & Krehenwinkel, H. (2020). High-throughput sequencing for community analysis: the promise of DNA barcoding to uncover diversity, relatedness, abundances and interactions in spider communities. Development Genes and Evolution, 230(2), 185–201. https://doi.org/10.1007/s00427-020-00652-x

Kinzelbach, R. (1992). The main features of the phylogeny and dispersal of the Zebra mussel Dreissena polymorpha. The Zebra Mussel Dreissena Polymorpha. Ecology, Biology Monitoring and First Applications in the Water Quality Management, 4, 5–17. Retrieved from Mollusc

Kozich, J. J., Westcott, S. L., Baxter, N. T., Highlander, S. K., & Schloss, P. D. (2013). Development of a dual-index sequencing strategy and curation pipeline for analyzing amplicon sequence data on the miseq illumina sequencing platform. Applied and Environmental Microbiology, 79(17), 5112–5120. https://doi.org/10.1128/AEM.01043-13

Koziol, A., Stat, M., Simpson, T., Jarman, S., DiBattista, J. D., Harvey, E. S., … Bunce, M. (2019). Environmental DNA metabarcoding studies are critically affected by substrate selection. Molecular Ecology Resources, 19(2), 366–376. https://doi.org/10.1111/1755-0998.12971

Kreeger, D. (1993). Seasonal patterns in utilization of dietary protein by the mussel Mytilus trossulus. Marine Ecology Progress Series, 95, 215–232. https://doi.org/10.3354/meps095215

Krehenwinkel, H., Kennedy, S., Pekár, S., & Gillespie, R. G. (2017). A cost-efficient and simple protocol to enrich prey DNA from extractions of predatory arthropods for large-scale gut content analysis by Illumina sequencing. Methods in Ecology and Evolution, 8(1), 126–134. https://doi.org/10.1111/2041-210X.12647

Krehenwinkel, H., Kennedy, S. R., Adams, S. A., Stephenson, G. T., Roy, K., & Gillespie, R. G. (2019). Multiplex PCR targeting lineage-specific SNPs: A highly efficient and simple approach to block out predator sequences in molecular gut content analysis. Methods in Ecology and Evolution, 10(7), 982–993. https://doi.org/10.1111/2041-210X.13183

Krehenwinkel, H., Kennedy, S. R., Rueda, A., Lam, A., & Gillespie, R. G. (2018). Scaling up DNA barcoding – Primer sets for simple and cost efficient arthropod systematics by multiplex PCR and Illumina amplicon sequencing. Methods in Ecology and Evolution, 9(11), 2181–2193. https://doi.org/10.1111/2041-210X.13064

Lavrentyev, P. J., Gardner, W. S., Cavaletto, J. F., & Beaver, J. R. (1995). Effects of the Zebra Mussel (Dreissena polymorpha Pallas) on Protozoa and Phytoplankton from Saginaw Bay, Lake Huron. Journal of Great Lakes Research, 21(4), 545–557. https://doi.org/10.1016/S0380-1330(95)71065-6

Leese, F., Sander, M., Buchner, D., Elbrecht, V., Haase, P., & Zizka, V. M. A. (2021). Improved freshwater macroinvertebrate detection from environmental DNA through minimized nontarget amplification. Environmental DNA, 3(1), 261–276. https://doi.org/10.1002/edn3.177

Lynggaard, C., Nielsen, M., Santos-Bay, L., Gastauer, M., Oliveira, G., & Bohmann, K. (2019). Vertebrate diversity revealed by metabarcoding of bulk arthropod samples from tropical forests. Environmental DNA, 1(4), 329–341. https://doi.org/10.1002/edn3.34

Machida, R. J., & Knowlton, N. (2012). PCR Primers for Metazoan Nuclear 18S and 28S Ribosomal DNA Sequences. PLoS ONE, 7(9), e46180. https://doi.org/10.1371/journal.pone.0046180

Macisaac, H. J., Lonnee, C. J., & Leach, J. H. (1995). Suppression of microzooplankton by zebra mussels: importance of mussel size. Freshwater Biology, 34(2), 379–387. https://doi.org/10.1111/j.1365-2427.1995.tb00896.x

Maguire, C. M., & Grey, J. (2006). Determination of zooplankton dietary shift following a zebra mussel invasion, as indicated by stable isotope analysis. Freshwater Biology, 51(7), 1310–1319. https://doi.org/10.1111/j.1365-2427.2006.01568.x

Mariani, S., Baillie, C., Colosimo, G., & Riesgo, A. (2019). Sponges as natural environmental DNA samplers. Current Biology, 29(11), R401–R402. https://doi.org/10.1016/j.cub.2019.04.031

Marquina, D., Esparza-Salas, R., Roslin, T., & Ronquist, F. (2019). Establishing insect community composition using metabarcoding of soil samples, and preservative ethanol and homogenate from Malaise trap catches: Surprising inconsistencies between methods. BioRxiv, 19(6), 1516–1530. https://doi.org/10.1101/597302

Matthews, J., Van der Velde, G., Bij de Vaate, A., Collas, F. P. L., Koopman, K. R., & Leuven, R. S. E. W. (2014). Rapid range expansion of the invasive quagga mussel in relation to zebra mussel presence in The Netherlands and Western Europe. Biological Invasions, 16(1), 23–42. https://doi.org/10.1007/s10530-013-0498-8

Miller, E. B., & Watzin, M. C. (2007). The effects of zebra mussels on the lower planktonic foodweb in Lake Champlain. Journal of Great Lakes Research, 33(2), 407–420. https://doi.org/10.3394/0380-1330(2007)33[407:TEOZMO]2.0.CO;2

Molloy, D. P., Karatayev, A. Y., Burlakova, L. E., Kurandina, D. P., & Laruelle, F. (1997). Natural enemies of zebra mussels: Predators, parasites, and ecological competitors. Reviews in Fisheries Science, 5(1), 27–97. https://doi.org/10.1080/10641269709388593

Newell, R. I. E. (2004). Ecosystem influences of natural and cultivated populations of suspension-feeding bivalve molluscs: A review. Journal of Shellfish Research, 23(1), 51–61.

Oksanen, J., Kindt, R., Legendre, P., O’Hara, B., Simpson, G. L., Solymos, P. M., … & Wagner, H. (2008). The vegan package. Community Ecology Package, 10(631–637), 190. Retrieved from https://bcrc.bio.umass.edu/biometry/images/8/85/Vegan.pdf

Paulus, M., Klein, R., & Teubner, D. (2018). Guideline for Sampling and Sample Processing Blue Mussel (Mytilus edulis complex). Environment Protection Agency (Ed.), Guidelines. Retrieved from https://www.umweltprobenbank.de/en/documents/publications/26658

Paulus, M., Teubner, D., Hochkirch, A., & Veith, M. (2014). Journey into the past: using cryogenically stored samples to reconstruct the invasion history of the quagga mussel (Dreissena rostriformis) in German river systems. Biological Invasions, 16(12), 2591–2597. https://doi.org/10.1007/s10530-014-0689-y

Pettersen, A. K., Turchini, G. M., Jahangard, S., Ingram, B. A., & Sherman, C. D. H. (2010). Effects of different dietary microalgae on survival, growth, settlement and fatty acid composition of blue mussel (Mytilus galloprovincialis) larvae. Aquaculture, 309(1–4), 115–124. https://doi.org/10.1016/j.aquaculture.2010.09.024

Richter, N. (2020). Entwicklung einer Next-Generation Sequencing basierten Methode zur Quantifizierung von taxonomischen Verunreinigungen in Deikantmuschelproben der Umweltprobenbank des Bundes. (Unpublished undergraduate thesis). University of Trier, Germany.

Schnell, I. B., Thomsen, P. F., Wilkinson, N., Rasmussen, M., Jensen, L. R. D., Willerslev, E., … Gilbert, M. T. P. (2012). Erratum: Screening mammal biodiversity using dna from leeches (Current Biology (2012) 22 (R262-R263)). Current Biology, 22(20), 1980. https://doi.org/10.1016/j.cub.2012.10.014

Seymour, M., Edwards, F. K., Cosby, B. J., Kelly, M. G., de Bruyn, M., Carvalho, G. R., & Creer, S. (2020). Executing multi-taxa eDNA ecological assessment via traditional metrics and interactive networks. Science of the Total Environment, 729, 138801. https://doi.org/10.1016/j.scitotenv.2020.138801

Sheppard, S. K., Bell, J., Sunderland, K. D., Fenlon, J., Skervin, D., & Symondson, W. O. C. (2005). Detection of secondary predation by PCR analyses of the gut contents of invertebrate generalist predators. Molecular Ecology, 14(14), 4461–4468. https://doi.org/10.1111/j.1365-294X.2005.02742.x

Shum, P., Barney, B. T., O’Leary, J. K., & Palumbi, S. R. (2019). Cobble community DNA as a tool to monitor patterns of biodiversity within kelp forest ecosystems. Molecular Ecology Resources, 19(6), 1470–1485. https://doi.org/10.1111/1755-0998.13067

Shumway, S. E., Davis, C., Downey, R., Karney, R., Kraeuter, J., Parsons, J., … Wikfors, G. (2003). Shellfish aquaculture — In praise of sustainable economies and environments. World Aquaculture, 34(4), 15–17.

Siegenthaler, A., Wangensteen, O. S., Soto, A. Z., Benvenuto, C., Corrigan, L., & Mariani, S. (2019). Metabarcoding of shrimp stomach content: Harnessing a natural sampler for fish biodiversity monitoring. Molecular Ecology Resources, 19(1), 206–220. https://doi.org/10.1111/1755-0998.12956

Sikder, M. M., Vestergård, M., Sapkota, R., Kyndt, T., & Nicolaisen, M. (2020). Evaluation of metabarcoding primers for analysis of soil nematode communities. Diversity, 12(10), 1–14. https://doi.org/10.3390/d12100388

Son, M. O. (2007). Native range of the zebra mussel and quagga mussel and new data on their invasions within the Ponto-Caspian Region. Aquatic Invasions, 2(3), 174–184. https://doi.org/10.3391/ai.2007.2.3.4

Sprung, M., & Rose, U. (1988). Influence of food size and food quantity on the feeding of the mussel Dreissena polymorpha. Oecologia, 77(4), 526–532. https://doi.org/10.1007/BF00377269

Stadhouders, R., Pas, S. D., Anber, J., Voermans, J., Mes, T. H. M., & Schutten, M. (2010). The effect of primer-template mismatches on the detection and quantification of nucleic acids using the 5′ nuclease assay. Journal of Molecular Diagnostics, 12(1), 109–117. https://doi.org/10.2353/jmoldx.2010.090035

Strohmeier T., Strand Ø., Alunno-Bruscia, M., Duinker, A., & Cranford, P. J. (2012). Variability in particle retention efficiency by the mussel Mytilus edulis. Journal of Experimental Marine Biology and Ecology, 412, 96–102. https://doi.org/10.1016/j.jembe.2011.11.006

Taberlet, P., Coissac, E., Pompanon, F., Brochmann, C., & Willerslev, E. (2012). Towards next-generation biodiversity assessment using DNA metabarcoding. Molecular Ecology, 21(8), 2045–2050. https://doi.org/10.1111/j.1365-294X.2012.05470.x

Team, Rs. (2015). RStudio: Integrated development for R. RStudio. RStudio, Inc., Boston, MA URL http://Www.Rstudio.Com, 42(42), 14. Retrieved from http://www.rstudio.com/.

Teubner, D., Klein, R., Tarricone, K., & Paulus, M. (2018). Guidelines for sampling, transport, storage and chemical characterization of environmental and human samples. Environment Protection Agency (Ed.): Guidelines for Sampling, Transport, Storage and Chemical Characterization of Environmental and Human Samples, Environment Protection Agency,Berlin. Retrieved from https://www.umweltprobenbank.de/en/documents/publications/26988%0A

Teubner, D., Wesslein, A. K., Rønne, P. B., Veith, M., Frings, C., & Paulus, M. (2016). Is a visuo-haptic differentiation of zebra mussel and quagga mussel based on a single external morphometric shell character possible? Aquatic Invasions, 11(2), 145–154. https://doi.org/10.3391/ai.2016.11.2.04

Thomsen, P. F., & Sigsgaard, E. E. (2019). Environmental DNA metabarcoding of wild flowers reveals diverse communities of terrestrial arthropods. Ecology and Evolution, 9(4), 1665–1679. https://doi.org/10.1002/ece3.4809

Turner, C. R., Barnes, M. A., Xu, C. C. Y., Jones, S. E., Jerde, C. L., & Lodge, D. M. (2014). Particle size distribution and optimal capture of aqueous macrobial eDNA. Methods in Ecology and Evolution, 5(7), 676–684. https://doi.org/10.1111/2041-210X.12206

Turon, M., Angulo-Preckler, C., Antich, A., Præbel, K., & Wangensteen, O. S. (2020). More Than Expected From Old Sponge Samples: A Natural Sampler DNA Metabarcoding Assessment of Marine Fish Diversity in Nha Trang Bay (Vietnam). Frontiers in Marine Science, 7. https://doi.org/10.3389/fmars.2020.605148

Valentin, R. E., Fonseca, D. M., Gable, S., Kyle, K. E., Hamilton, G. C., Nielsen, A. L., & Lockwood, J. L. (2020). Moving eDNA surveys onto land: Strategies for active eDNA aggregation to detect invasive forest insects. Molecular Ecology Resources, 20(3), 746–755. https://doi.org/10.1111/1755-0998.13151

van der Heyde, M., Bunce, M., Wardell-Johnson, G., Fernandes, K., White, N. E., & Nevill, P. (2020). Testing multiple substrates for terrestrial biodiversity monitoring using environmental DNA metabarcoding. Molecular Ecology Resources, 20(3), 732–745. https://doi.org/10.1111/1755-0998.13148

Vaughn, C. C., Nichols, S. J., & Spooner, D. E. (2008). Community and foodweb ecology of freshwater mussels. Journal of the North American Benthological Society, 27(2), 409–423. https://doi.org/10.1899/07-058.1

Vestheim, H., & Jarman, S. N. (2008). Blocking primers to enhance PCR amplification of rare sequences in mixed samples -A case study on prey DNA in Antarctic krill stomachs. Frontiers in Zoology, 5(1), 1–11. https://doi.org/10.1186/1742-9994-5-12

Wagner, G., Bartel, M., Klein, R., Neitzke, M., Nentwich, K., Paulus, M., & Quack, M. (2003). Guideline for Sampling and Sample Treatment: Zebra Mussel (Dreissena polymorpha). Umweltprobenamt Und Umweltbundesamt, 1–15.

Wai, H. W., & Levinton, J. S. (2004). Culture of the blue mussel Mytilus edulis (Linnaeus, 1758) fed both phytoplankton and zooplankton: A microcosm experiment. Aquaculture Research, 35(10), 965–969. https://doi.org/10.1111/j.1365-2109.2004.01107.x

Wickham, H., Chang, W., Henry, L., Pedersen, T. L., Takahashi, K., Wilke, C., … Dunnington, D. (2016). ggplot2: Elegant Graphics for Data Analysis. New York: Springer.

Wilcox, T. M., McKelvey, K. S., Young, M. K., Lowe, W. H., & Schwartz, M. K. (2015). Environmental DNA particle size distribution from Brook Trout (Salvelinus fontinalis). Conservation Genetics Resources, 7(3), 639–641. https://doi.org/10.1007/s12686-015-0465-z

Zhang, J., Kobert, K., Flouri, T., & Stamatakis, A. (2014). PEAR: A fast and accurate Illumina Paired-End reAd mergeR. Bioinformatics, 30(5), 614–620. https://doi.org/10.1093/bioinformatics/btt593

