## Supplement for "Molecular diet analysis in zebra and quagga mussels (*Dreissena* spp.) and an assessment of the utility of aquatic filter feeders as biological eDNA filters"

### Appendix

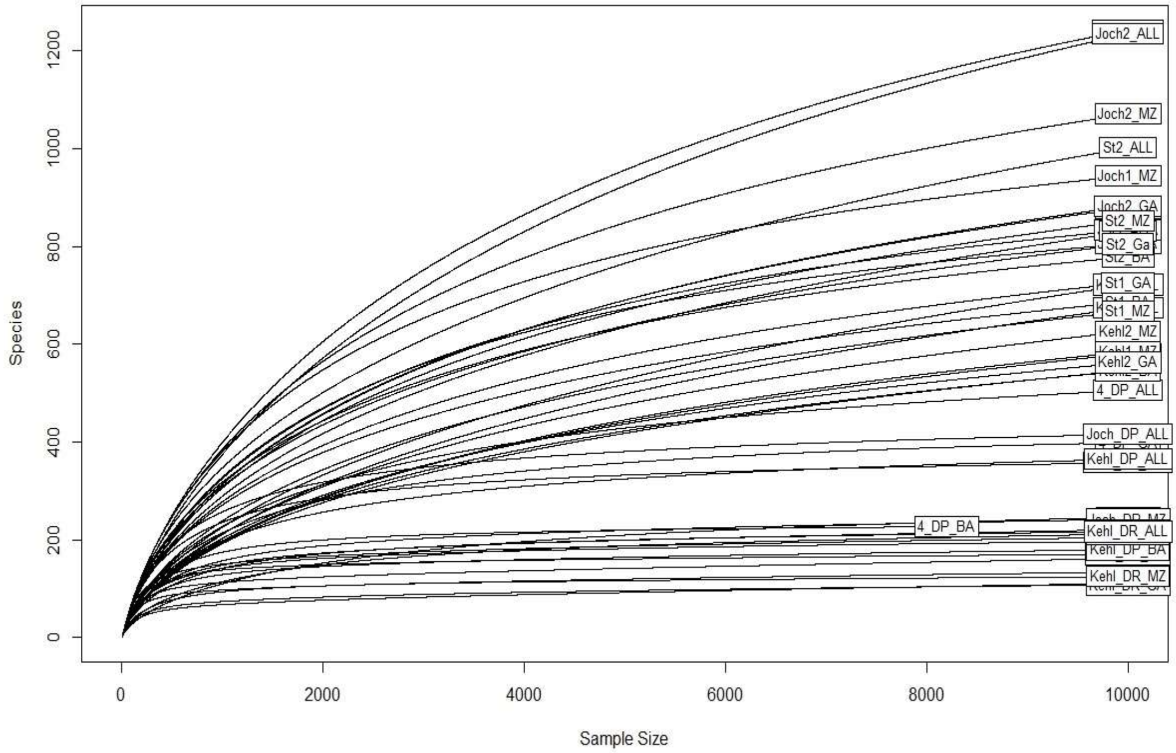


*Suppl. Figure 1 Rarefaction curves for all samples showing a saturation of diversity at aroound 10,000 reads.*

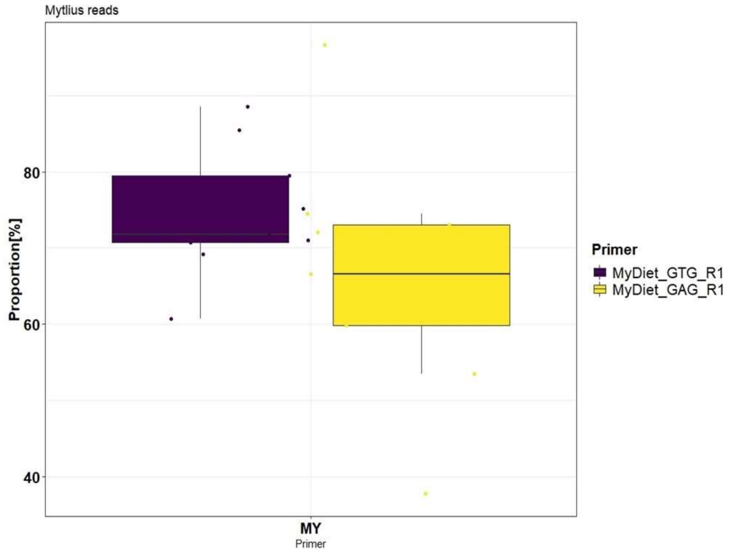


*Suppl. Figure 2) a) Recovered proportion of blue mussel reads for the two general metazoan blocking primer combinations.*

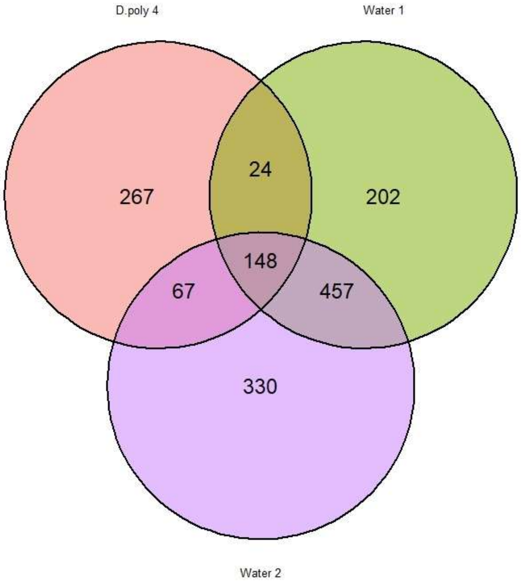


*Suppl. Figure 3 Venn diagram showing the recovered OTU numbers and their overlap between* D. polymorpha *and the two eDNA water samples for the site at Lake Stechlin.*

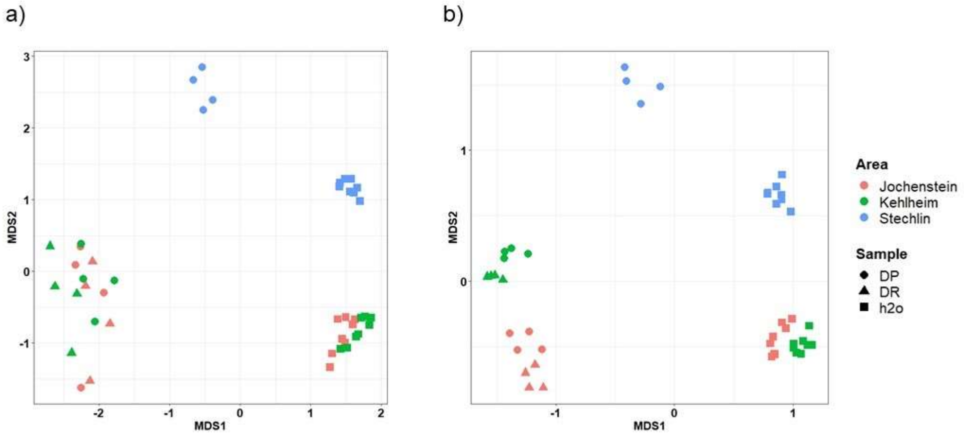


*Suppl. Figure 4 NMDS plots showing community differentiation of eDNA water samples and* Dreissena *mussel samples for the two sites at the Danube (Kelheim and Jochenstein) and the Stechlin lake. The first axis separates water (left) and mussels (right). All samples are well separated by site, with water and mussel samples showing similar patterns of differentiation. The shape distinguishes water samples as well as* D. polymorpha *(DP) and* D. rostriformis *(DR). a) Shows the NMDS plot (stress = 0.088), with commensal and parasite OTUs of the mussels removed, b) (stress = 0.089) shows the same plot including parasites and commensals.*

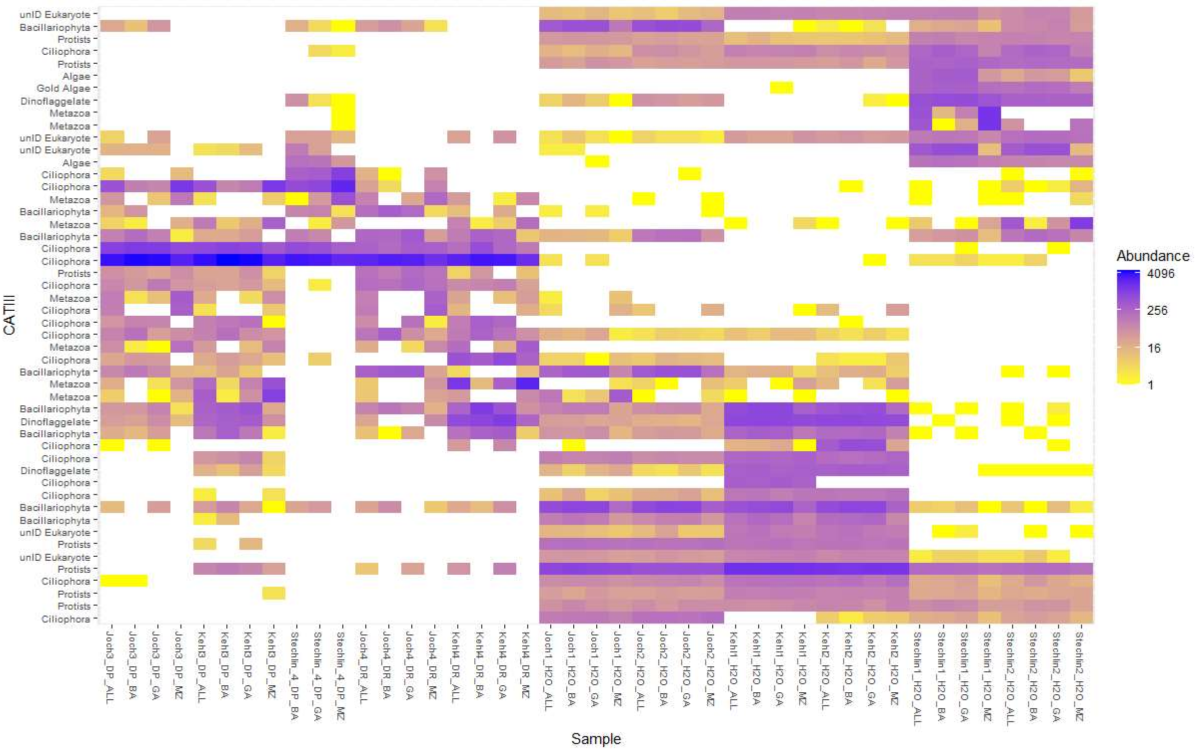


*Suppl. Figure 5 Heatmap of the 50 most abundant OTUs in the data set showing a clear differentiation of taxonomic composition between mussel samples (left) and water samples (right). As shown in the NMDS, there is no evidence that the primer affects the overall composition drastically. At the same time, each sampling site has a characteristic OTU composition despite the fact that all two of the three sites (Jochenstein, Kelheim) are situated on the same river (Danube).*

Table 1: 18S primer combinations used in this study. All primers were designed and tested for 18SrDNA amplification in *Dreissena polymorpha*, *Dreissena rostriformis, Mytilus edilus* and environmental DNA water samples. The table shows the recommended combination of forward and reverse primers. The taxon specific sequences, used to block mussels are highlighted in bold within the primer sequence. The corresponding sequence to the blocking nucleotides in *Dreissena* are shown in the "*Dreissena* sequence" column. The "Reduction" column represents the proportion of blocked Mussel reads (*Dreissena* spp. and *Mytilus edilus*. (*only tested *in silico*)

| **Name** | **Forward (5'- 3')** | **Name** | **Reverse (5'- 3')** | **Dreissena sequence** | **Reduction [%]** |
|---|---|---|---|---|---|
| EukF1 | CAATAACAGGTCTGTGATGC | DreissDiet_TAA_R | **AA**G**TAA**AAGTCGTAACAAGGTTTYCG | TC/TAA | 82.11 ± 6.36 |
| NonMetazoaF1 | CTGTGATGCCCTTAGATGTYC**T** | DreissDiet_GTG_R | **AA**G**GTG**AAGTCGTAACAAGGTTTC | G/TC/TAA | 99.9 ±0.37 |
| NonMetazoaF1 | CTGTGATGCCCTTAGATGTYC**T** | DreissDiet_GAG_R | **AA**G**GAG**AAGTCGTAACAAGGTYTC | G/TC/TAA | 99.9 ±0.006 |
| NonMetazoaF1 | CTGTGATGCCCTTAGATGTYC**T** | NonMetazoaDiet_GTG_R1 | **GTG**AAGTCGTAACAAGGTTTCCG | G/TAA | 25.55 ± 8.63 |
| NonMetazoaF1 | CTGTGATGCCCTTAGATGTYC**T** | NonMetazoaDiet_GAG_R1 | **GAG**AAGTCGTAACAAGGTYTCCG | G/TAA | 33.91 ± 16.28 |
| EukF2 | GTCCCTGCCCTTTGTACA | DreissDiet_TAA_R | **AA**G**TAA**AAGTCGTAACAAGGTTTYCG | TC/TAA | * |
| EukF2 | GTCCCTGCCCTTTGTACA | DreissDiet_GTG_R | **AA**G**GTG**AAGTCGTAACAAGGTTTC | TC/TAA | * |
| EukF2 | GTCCCTGCCCTTTGTACA | DreissDiet_GAG_R | **AA**G**GAG**AAGTCGTAACAAGGTYTC | TC/TAA | * |
| EukF2 | GTCCCTGCCCTTTGTACA | NonMetazoaDiet_GTG_R1 | **GTG**AAGTCGTAACAAGGTTTCCG | TAA | * |
| EukF2 | GTCCCTGCCCTTTGTACA | NonMetazoaDiet_GAG_R1 | **GAG**AAGTCGTAACAAGGTYTCCG | TAA | * |
